# Temporal signal and the phylodynamic threshold of SARS-CoV-2

**DOI:** 10.1101/2020.05.04.077735

**Authors:** Sebastian Duchene, Leo Featherstone, Melina Haritopoulou-Sinanidou, Andrew Rambaut, Philippe Lemey, Guy Baele

## Abstract

The ongoing SARS-CoV-2 outbreak marks the first time that large amounts of genome sequence data have been generated and made publicly available in near real-time. Early analyses of these data revealed low sequence variation, a finding that is consistent with a recently emerging outbreak, but which raises the question of whether such data are sufficiently informative for phylogenetic inferences of evolutionary rates and time scales. The phylodynamic threshold is a key concept that refers to the point in time at which sufficient molecular evolutionary change has accumulated in available genome samples to obtain robust phylodynamic estimates. For example, before the phylodynamic threshold is reached, genomic variation is so low that even large amounts of genome sequences may be insufficient to estimate the virus’s evolutionary rate and the time scale of an outbreak. We collected genome sequences of SARS-CoV-2 from public databases at 8 different points in time and conducted a range of tests of temporal signal to determine if and when the phylodynamic threshold was reached, and the range of inferences that could be reliably drawn from these data. Our results indicate that by February 2^nd^ 2020, estimates of evolutionary rates and time scales had become possible. Analyses of subsequent data sets, that included between 47 to 122 genomes, converged at an evolutionary rate of about 1.1×10^−3^ subs/site/year and a time of origin of around late November 2019. Our study provides guidelines to assess the phylodynamic threshold and demonstrates that establishing this threshold constitutes a fundamental step for understanding the power and limitations of early data in outbreak genome surveillance.

## Main text

Pathogen genome sequence data are increasingly recognised as a key asset in outbreak investigations. Phylodynamic analyses of these data can be used to infer the time and location of origin of an outbreak, the viral evolutionary rate, epidemiological dynamics, and demographic patterns (du Plessis and Stadler 2015; Baele et al. 2017). These inferences, however, rely on the genome data being sufficiently informative.

The ongoing novel coronavirus outbreak (SARS-CoV-2) marks the first time that genome sequence data have been generated and shared publicly as soon as the virus started spreading. The time of origin of SARS-CoV-2 is a pressing question at early stages of the outbreak because it impacts our understanding of its spread and emergence. In practice, the sampling times of genomes can be used to calibrate the molecular clock and infer the viral evolutionary rate and the timescale of the outbreak (Korber et al. 2000). The underlying assumption is that molecular evolution occurs at a predictable rate over time and that the sampling window is sufficiently wide as to capture a measurable amount of evolutionary change in the sampled genomes. Under the condition that the sampling window is sufficiently wide and the evolutionary rate sufficiently high, and genome sequences long enough, the data can be treated as having been obtained from a measurably evolving population (Drummond et al. 2003; Biek et al. 2015). If this is not the case, the data are considered to have no temporal signal and any estimates from the molecular clock are therefore spurious (Duchêne et al. 2015; Murray et al. 2015).

The term ‘phylodynamic threshold’ pertains to the question of whether a virus has had sufficient time to evolve since its origin so as to warrant tip-dating calibration, under the assumption that genome data from early stages of the outbreak are available (Hedge et al. 2013). Therefore, applying statistical tests of temporal signal to genome data as they are collected can reveal when the phylodynamic threshold is reached. Such analyses are essential to determine the limitations of genome data and the range of inferences that can be reliably drawn from them over time.

Root-to-tip regression is typically used as an informal assessment of temporal signal (Rambaut et al. 2016). While not a statistical test, it is however a valuable visual tool of clocklike behaviour and of outlier detection (e.g. due to mislabelling, contamination or sequencing errors). Root-to-tip regression consists of estimating an unrooted phylogenetic tree with branch lengths in units of substitutions per site and conducting a regression of the distance from the root to each of the tips as a function of their sampling times (Gojobori et al. 1990; Drummond et al. 2003). Under clocklike evolution and with a wide sampling window, the slope corresponds to a crude estimate of the evolutionary rate, the intercept with the time axis represents the time of origin, and the coefficient of determination, R^2^, may reflect the degree of clocklike behaviour.

Formal approaches to assess temporal signal include date-randomisation tests and Bayesian evaluation of temporal signal (BETS) (Duchêne et al. 2015; Murray et al. 2015; Duchene et al. 2019). Date randomisation tests consist of repeating the analysis several times with permuted sampling times to generate a ‘null’ distribution of evolutionary rate estimates. The data are considered to have temporal signal if the estimate obtained with the correct sampling times does not overlap with those of the randomisations. In contrast, BETS consists of comparing the statistical fit of models that include the correct sampling times, no sampling times, or permuted sampling times. The premise of BETS is that if the data have temporal signal, using the correct sampling times should have the highest statistical fit (Duchene et al. 2019). For example, if the sampling window over which the genome data have been collected is very short, such that the data have no temporal signal, then the sampling times are not meaningful and a model incorporating the correct sampling times may not have an improved statistical fit over a model that ignores differences in sampling times. In contrast, if the sampling window is wide enough as to capture many substitutions, using the correct sampling times is expected to result in higher model fit than using permuted sampling times or no sampling times. In a Bayesian context, model fit is determined through the marginal likelihood, and a model is preferred over another according to their ratio of marginal likelihoods, known as the Bayes factor (Kass and Raftery 1995). Marginal likelihoods are typically reported on a logarithmic scale, where a log Bayes factors of at least 1 is considered as positive evidence in favour of a model.

### Results

We collected SARS-CoV-2 genome data from the Global Initiative on Sharing All Influenza Data (GISAID) and from GenBank at 8 time points from January 23^rd^ to February 24^th^ 2020 (Table 1). Thus, each time point represents a ‘snapshot’ of the genome data available to that date. Our data only included genomic sequences from human samples, with sequence lengths of at least 28,000 nucleotides and, with high coverage as determined in GISAID (see supplementary material Table S1 for accession numbers). To minimise the impact of potential sequencing errors in our alignments, we deleted obvious errors upon visual inspection and compared our phylogenetic trees to those obtained by other groups (virological.org) and those from the Nextstrain workflow (Hadfield et al. 2018).

**Table 1.** Description of data snapshots of SARS-CoV-2.

| Publication date range<br>(from Jan 10 2020) | Number of<br>genomes | Sampling window (from<br>Dec 23 2019) |
| --- | --- | --- |
| Jan 23 | 22 | Jan 17 2020 |
| Feb 2 | 47 | Jan 27 2020 |
| Feb 6 | 55 | Jan 28 2020 |
| Feb 10 | 66 | Feb 3 2020 |
| Feb 15 | 90 | Feb 7 2020 |
| Feb 18 | 95 | Feb 9 2020 |
| Feb 21 | 109 | Feb 9 2020 |
| Feb 24 | 122 | Feb 10 2020 |

We conducted Bayesian phylogenetic analyses using BEAST v1.10 using two molecular clock models; a strict clock (SC) and an uncorrelated relaxed clock with an underlying lognormal distribution (UCLN). We set an exponential growth coalescent tree prior, which is appropriate for the early stages of an outbreak and which has been recently used to infer the basic reproductive number and growth rate of SARS-CoV-2 (Volz et al. 2020). For our model comparison in BETS we estimated (log) marginal likelihoods using generalised stepping-stone sampling (Fan et al. 2011; Baele et al. 2016).

Our BETS analyses provided evidence against significant temporal signal in the genome data available up to January 23^rd^ 2020 (n=22 genomes). In this data set, the highest model fit to the data was found for analyses with permuted sampling times, followed by those with no sampling times (Figure 1). The evidence for models with no sampling times was very strong, with log Bayes factors of 7.5 for the best model with no sampling times relative to those without sampling times. All data sets obtained subsequently, from February 2^nd^ with at least 47 genomes supported the inclusion of the correct sampling times, with log Bayes factors of at least 20 for models with the correct sampling times over those without sampling times. The log Bayes factors for the models with correct sampling times over those with permuted sampling times were at least 5, which is considered as very strong evidence in favour of temporal signal (Kass and Raftery, 1995).

**Figure 1.**
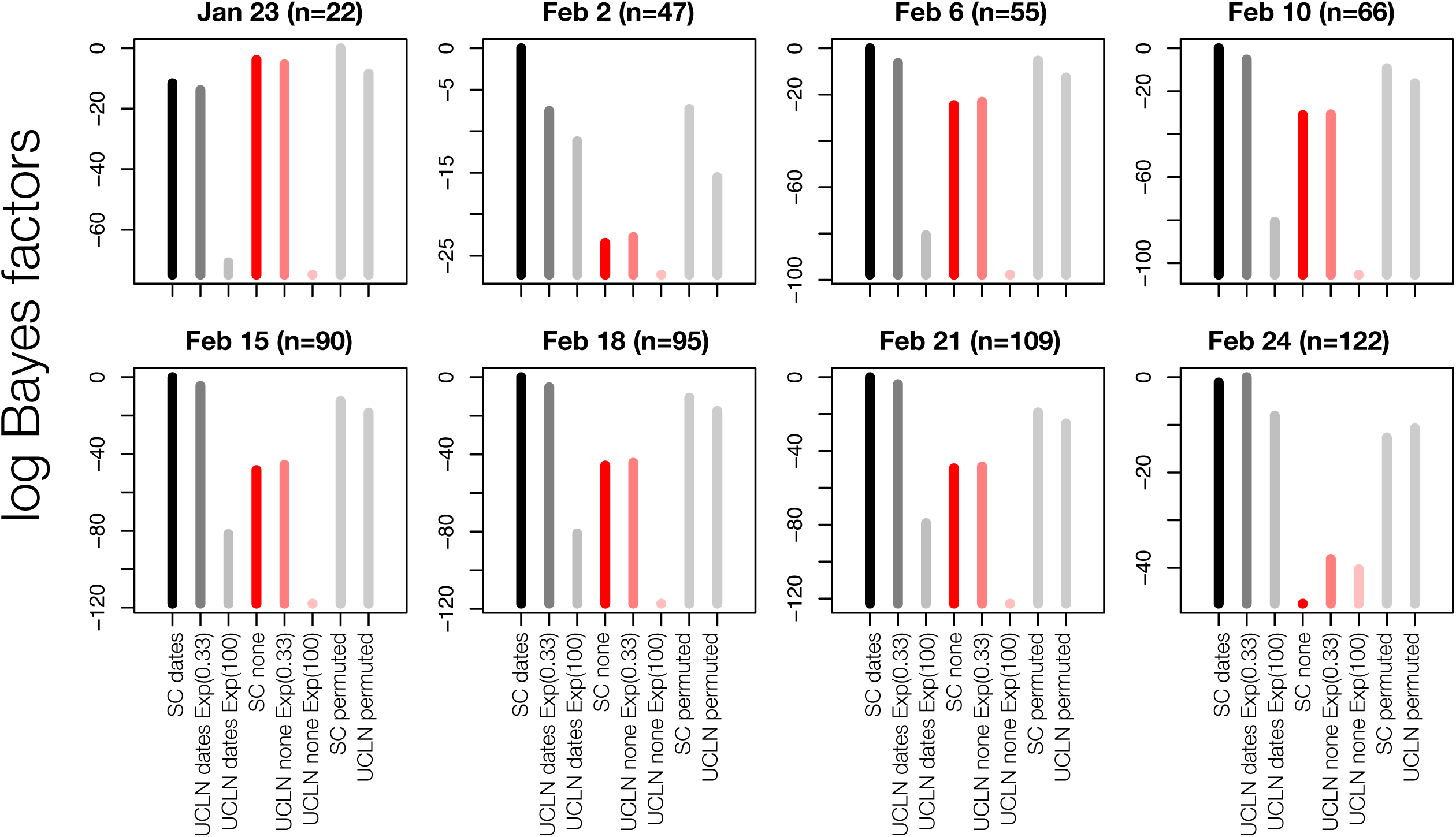
Bayesian evaluation of temporal signal (BETS) results. Each panel corresponds to a snapshot data set collected up to a given month and day in 2020 and with a certain number, n, of genomes. The *y*-axis represent the log Bayes factors, where the best-performing model has a value of 0. Each bar corresponds to an analysis configuration for BETS, with two possible molecular clock models: the strict (SC) and the uncorrelated relaxed clock with an underlying lognormal distribution (UCLN). For the UCLN, we considered two possible priors on the standard deviation of the lognormal distribution: an exponential distribution with mean 0.33 or with mean 100, labelled as Exp(0.33) and Exp(100), respectively. The sampling times could be configured using the true values (dates), no sampling times (none), or permuted, with these latter two options indicating no temporal signal. Black and dark grey bars correspond to analyses with the correct sampling times with the SC or UCLN clock models, respectively. Dark and light red bars are for analyses with no sampling times with these two clock models, and all light grey bars are for analyses with permuted sampling times.

For comparison, we also conducted root-to-tip regressions for the eight snapshot data sets (Figure 2). The R^2^ values ranged between 0.11 and 0.2. We did not find an association between R^2^ and the number of genome samples included. This result may stand in contrast to the expectation that including more independent data should reduce the effect of stochasticity, but the data sets here have an inherently high degree of non-independence. The slopes of the regressions ranged from 6.7×10^−4^ to 8.8×10^−4^ subs/site/year and the intercept with the X-axis (i.e. the time to the most recent common ancestor) from 2019.83 to 2019.86.

**Figure 2.**
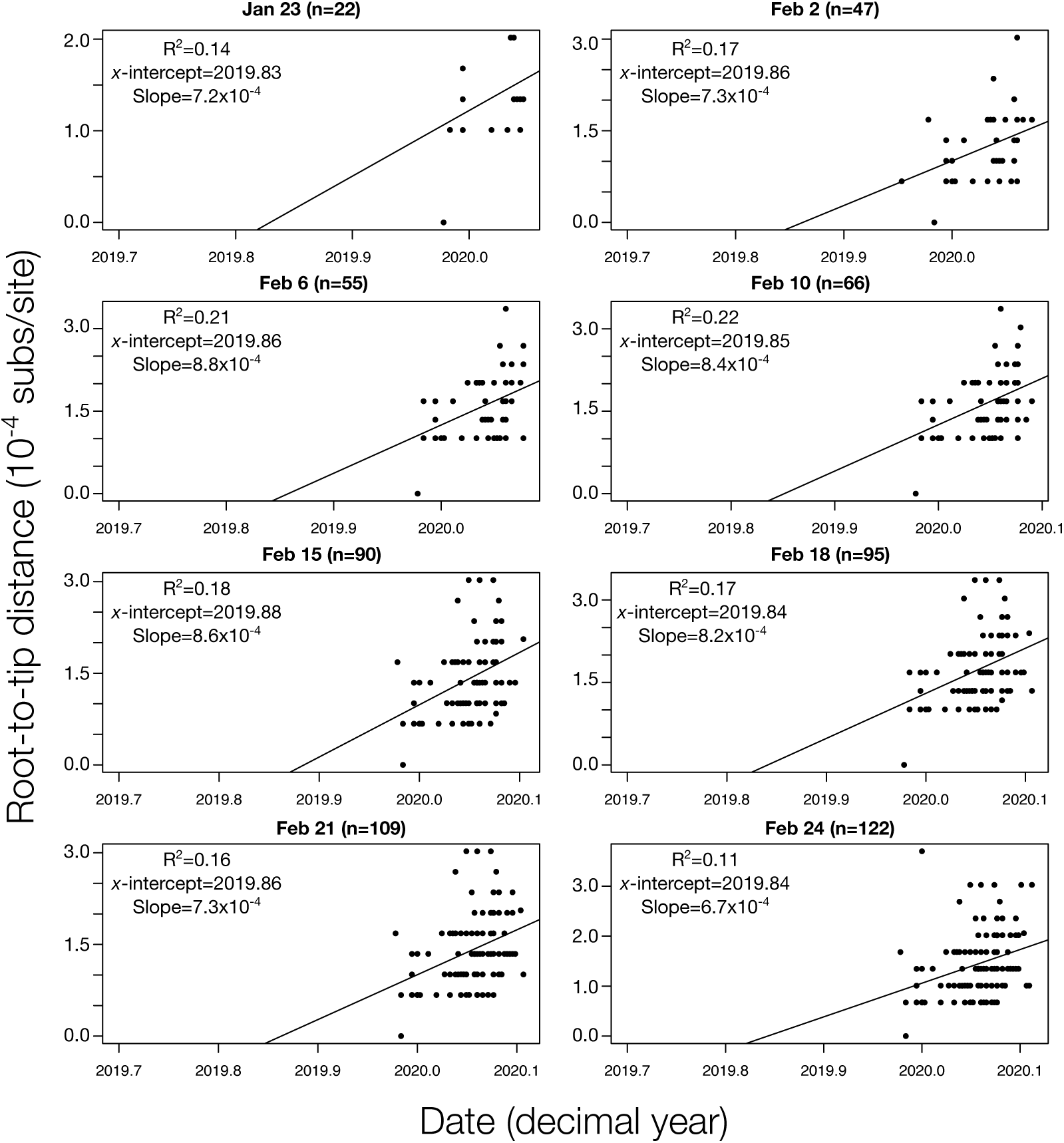
Root-to-tip regressions for snapshot data sets. The *y*-axis corresponds to the root-to-tip distance of phylogenetic trees with branch lengths in units of substitutions per site. The *x*-axis represents calendar time. Each point corresponds to a tip in the tree. The regression line is the best fitting line using the root position that maximised R^2^. The R^2^, the intercept with the *x*-axis (*x*-intercept), and slope are shown for each data set, with the latter two representing crude estimates of the evolutionary rate and time of origin, respectively.

Interestingly, all data sets with temporal signal favoured the SC over the UCLN model with the exception of that collected up to February 24^th^, with 122 genomes, where the log Bayes factor of the UCLN over the SC was 1.81 (Figure 1). The fact that the SC had high support in data sets collected prior to February 24^th^ probably indicates that they may not be sufficiently informative as to warrant modelling evolutionary rate variation across branches through the UCLN, rather than evidence of strict clocklike behaviour.

A potential reason for why the SC is favoured over the UCLN in many cases is that the default prior on the standard deviation of the lognormal distribution of the UCLN is an exponential distribution with mean 0.33, that has a high density at 0, corresponding to a very low amount of among-lineage rate variation. Intuitively, if the data have low information content, the prior may have a strong influence on the posterior, relative to the data, such that the posterior for this parameter might also be concentrated on 0. In this case, the UCLN may appear overparameterised and the SC would be favoured. We investigated the robustness of model selection to the prior on this parameter repeating the UCLN analyses with an exponential distribution with mean 100 as the prior for this parameter. Using this less informative prior consistently resulted in a worse model fit across all data sets, and thus did not affect our assessment of temporal signal.

If we restrict our attention to the UCLN with the less informative prior for the January 23^rd^ data set, the model that includes sampling times is favoured over that with no sampling times, with a log Bayes factor of 17. If one ignored all other models and priors, this result would indicate the presence of temporal signal. This finding stands in contrast to the SC and UCLN with the more informative prior, which have much higher model fit (Figure1; Supplementary material Table S2). Consequently, assessing temporal signal using BETS should involve comparing a range of clock models and careful consideration of the prior on their respective parameters (Duchene et al. 2019).

We also considered comparisons of prior and posterior distributions to assess the extent to which the data were informative about particular parameters. Our expectation is that the posterior should have a lower variance relative to the prior as more data are included. We considered our estimates of the growth rate (*r*) and scaled population size (*Φ*) of the exponential coalescent tree prior, the virus’s evolutionary rate and the time of origin of the outbreak. An important consideration here is that our method of inspecting the prior consists in running the analyses with no sequence data. Thus, the resulting distributions represent the ‘effective’, rather than the ‘marginal’ (i.e. user-specified) prior. The effective prior is the prior conditioned on the number of samples and their ages, the coalescent process and their interaction, whereas the marginal prior is the actual distribution that one sets in the program. In practice, the effective and marginal prior sometimes differ for parameters that pertain to the tree prior (Warnock et al. 2012; Boskova et al. 2018).

Although our marginal priors are identical for all snapshot datasets, we noted that the effective prior differed between data sets for *r, Φ*, and the time of origin (Figure 3). The posterior from the January 23^*rd*^ snapshot, with 22 genomes, was very uncertain for all parameters. For example, the time of origin using the SC ranged from late 2018 to early December 2019. The posterior for *Φ* was also more uncertain than its effective prior, which coincides with high uncertainty in the rate and the time of origin.

**Figure 3.**
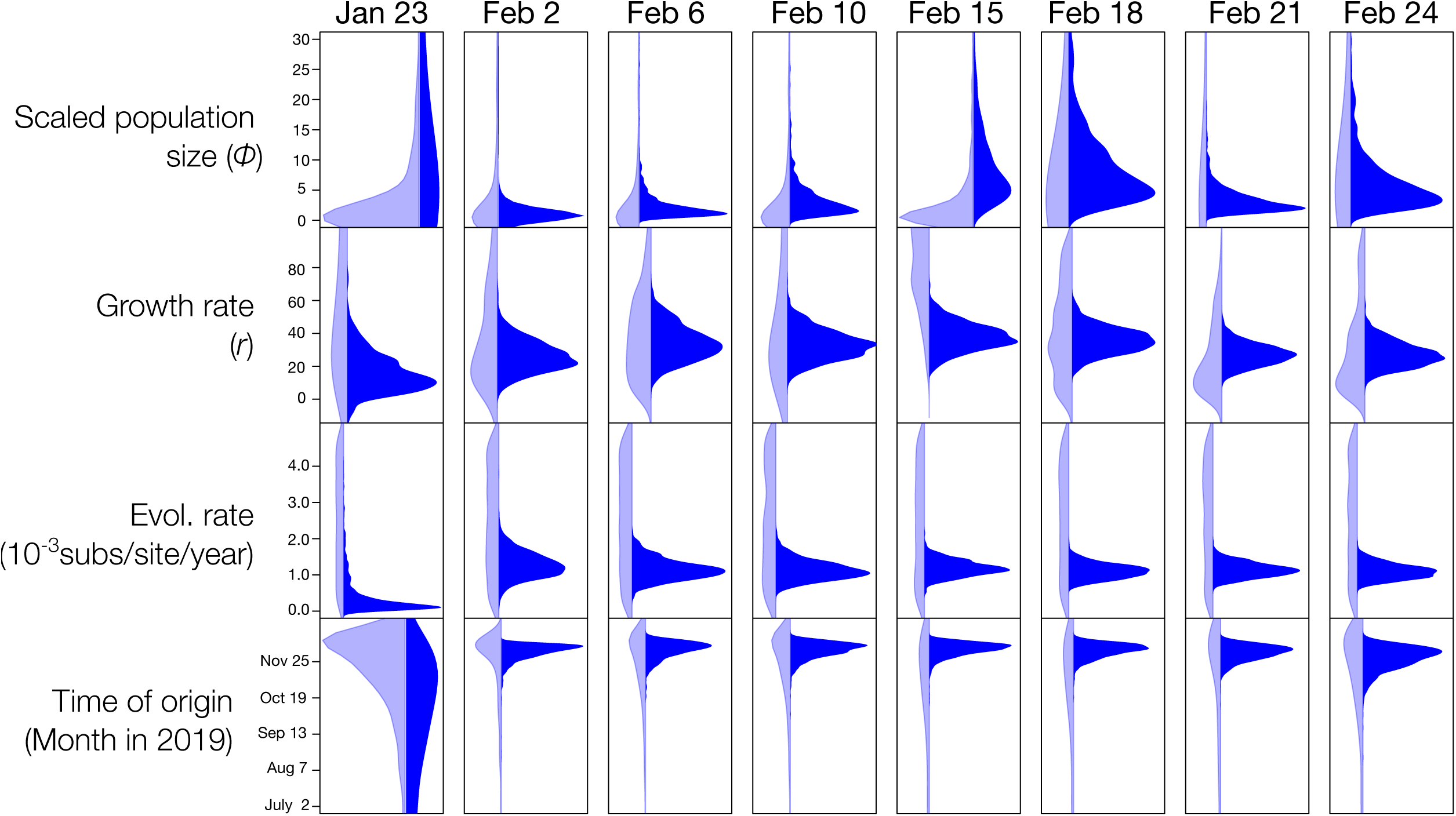
Prior and posterior densities for parameters of interest using the molecular clock model with best fit for all snapshot data set (SC for all data sets, except for February 24^th^, where the UCLN was chosen). The *y*-axis corresponds to parameter values, while the x-axis represents the relative density. Light blue densities correspond to the effective prior, while those in dark blue show the posterior.

Our snapshot data sets collected from February 2^nd^, with at least 47 genome samples, yielded posterior distributions that were much narrower than their respective effective priors and those of the January 23^rd^ snapshot. Our estimates of the evolutionary rate from February 2^nd^ converged at a mean of around 1.1×10^−3^ substitutions per site per year. The uncertainty in this parameter for the largest data set (February 24^th^, with 122 genomes) using the UCLN clock model is reflected by a 95% credible interval (CI) of between 7.03×10^−4^ and 1.5×10^−3^ substitutions per site per year. Similarly, the time of origin converged to a mean of late November 2019 and with a 95% CI for the February 24^th^ data set of between late October to mid-December 2019.

Posterior estimates for parameters *r* and *Φ*, differed substantially from their effective priors, although not to the extent that the evolutionary rate and the time of origin did. In particular, the posterior of the time of origin is several times narrower than the prior in all data sets from February 2^nd^, whereas the posterior for *r* in the largest data set (February 24^th^) is only about 2 times narrower than its respective effective prior (Figure 3). Our estimates of *r* and *Φ* did not converge between snapshot data sets, as was the case for the evolutionary rate and time of origin. However, we do not necessarily expect this to happen. For instance, *Φ* is proportional to the number of infected individuals at the time of collection of the latest sample (Wallinga and Lipsitch 2007; Boskova et al. 2014), which is expected to increase as the outbreak progresses. Similarly, *r* is proportional to the reproductive number *R*_***e***_, (i.e. the average number of secondary infections), is expected to decline over time as the number of susceptible individuals decreases and is expected to be affected by growing spatial structure.

### Discussion

The question of whether a viral outbreak has attained the phylodynamic threshold is a highly relevant concept for emerging outbreaks, because it is informative about the amount of sequence data, their temporal spread, and how much evolutionary change has accumulated in the viral genome. The phylodynamic threshold requires a strong assumption about the evolutionary rate based on closely related viruses, and it can be understood as the point in time when sequence data are sufficiently informative about the evolutionary dynamics that shape an outbreak, i.e. when the population is measurably evolving. The routine application of tests of temporal signal can effectively answer this question in nearly real-time. Our application of BETS (Duchene et al., 2019) to data snapshots from the early stages of the outbreak revealed that the phylodynamic threshold of SARS-CoV-2 was reached by about February 2^nd^, when 47 genomes were available sampled over 35 days.

Our finding that the phylodynamic threshold was attained within about two months of the estimated start of the outbreak demonstrates that Bayesian phylodynamic approaches can capitalise on early collected genome data to make inferences about evolutionary processes, particularly the viral evolutionary rate and the outbreak’s time of origin. Our estimates of these two parameters were consistent after the phylodynamic threshold was reached, and also matched previous estimates posted on virological.org and elsewhere (Taiaroa et al. 2020; Volz et al. 2020). Increasing the number of sequences leads to more precise estimates of the evolutionary rate, but we found only marginal improvements in precision after 109 sequences (February 21^st^). The SC was preferred over the UCLN in most data sets. The fact that the UCLN was only supported after 122 sequences were included suggests that the statistical power necessary to support such a relaxed clock model may require more informative data than those available at the early stages in the outbreak. We anticipate that the UCLN will be favoured over the SC in analyses of larger data sets of SARS-CoV-2.

A key consideration concerning the presence of temporal signal in the data is that this does not necessarily imply that demographic parameters can be reliably estimated using genome sequence data. Comparing the effective prior and posterior is important to assess the information content of the data, but it is not an assessment of the reliability of the estimates. For example, *Φ* is generally inversely correlated with the root height, such that if the data have temporal signal, the prior and posterior for this parameter will substantially differ. However, this parameter is proportional to the number of infected individuals at present under the assumption that the number of infections grows exponentially in a deterministic fashion and in the absence of population structure. Clearly, the extent to which the data meet these conditions can affect the interpretation and reliability of such epidemiological parameters. More realistic tree priors may be warranted here, such as those that account for population structure and the sampling process (Scire et al. 2020). In sum, whether the phylodynamic threshold coincides with reliability in estimates of epidemiological parameters depends on the information content in the data, but also on the tree prior and its underlying assumptions.

Ongoing analyses of SARS-CoV-2 will reveal important aspects regarding its evolutionary origin and epidemiological dynamics. On a global scale, the virus is well beyond its phylodynamic threshold, but tests of temporal signal, as applied here, will still be key to understand the timescale of local transmission.

### Methods

We downloaded genome sequence data from GISAID or GenBank, and aligned them using MAFFT (Katoh et al. 2002). We curated the data through comparison with data sets available at virological.org and visual inspections of our alignments (Supplementary Materials, Table S1). We only included sequences from humans, that were at least 28,000 nucleotides long, and with high coverage.

### Bayesian phylogenetic analyses

We analysed each data snapshot in BEAST (Suchard et al. 2018) using the HKY+Γ substitution model. We set a Markov chain Monte Carlo (MCMC) length of 10^7^ steps, sampling every 10^3^ steps. We determined sufficient sampling by verifying that the effective sample size of key parameters was at least 200 using Tracer v1.7 (Rambaut et al. 2018). We assessed temporal signal using BETS (Duchene et al. 2019). We compared the statistical fit of two molecular clock models, SC and UCLN, and three configurations of sampling times; the correct sampling times, no sampling times, and permuted sampling times, with the latter two corresponding to a lack of temporal signal. For each combination of molecular clock model and sampling times we calculated the (log) marginal likelihood using generalised stepping-stone sampling (Baele et al. 2016), for which we employed 200 path steps with a chain length for each power posterior of 10^5^ iterations. We chose priors for all parameters that respected their respective domains, but that were not overly informative, and all of which are proper (i.e. the area under the curve is 1.0; (Baele et al. 2013)) (Table 2). According to BETS, a data set is considered to have temporal signal if (log) Bayes factors support a model with the correct sampling times (Duchene et al. 2019).

**Table 2.**
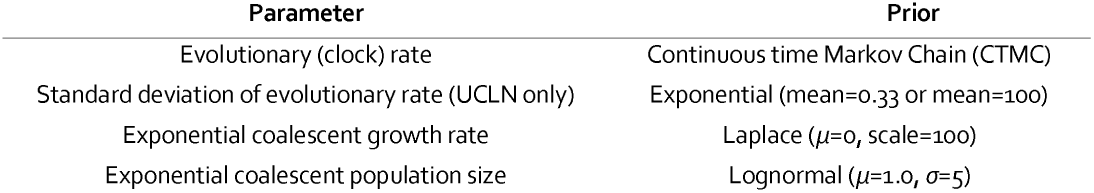
Prior distributions used for key parameters.

| Parameter | Prior |
| --- | --- |
| Evolutionary (clock) rate | Continuous time Markov Chain (CTMC) |
| Standard deviation of evolutionary rate (UCLN only) | Exponential (mean=0.33 or mean=100) |
| Exponential coalescent growth rate | Laplace ($\mu=0$, scale=100) |
| Exponential coalescent population size | Lognormal ($\mu=1.0$, $\sigma=5$) |

Our comparison of the prior and posterior distributions of key parameters require obtaining the effective, rather than the marginal prior. The effective prior can be obtained by running the analysis in BEAST with no sequence data, which is equivalent to ignoring the sequence likelihood and is done by selecting the option ‘sample from prior’ in BEAUti, the graphical interface accompanying the BEAST software package (Suchard et al. 2018). All Bayesian phylogenetic analyses were conducted on the SPARTAN high-performance computing service of the University of Melbourne (Meade et al. 2017).

### Root-to-tip regression

We estimated phylogenetic trees using maximum likelihood inference as implemented in IQ-tree v1.6 (Minh et al. 2020), with the optimal substitution model determined by the program. We used these trees to obtain root-to-tip regressions in TempEst v1.5 (Rambaut et al. 2016) by selecting the root position that maximised R^2^.

## Acknowledgements

We thank all those who have contributed sequences to the GISAID database (https://www.gisaid.org/). SD was supported by an Australian Research Council Discovery Early Career Researcher Award (DE190100805) and an Australian National Health and Medical Research Council grant (APP1157586). A.R. is supported by the Wellcome Trust (Collaborators Award 206298/Z/17/Z―ARTIC network). PL acknowledges funding from the European Research Council under the European Union’s Horizon 2020 research and innovation programme (grant agreement no. 725422-ReservoirDOCS) and the Research Foundation -- Flanders (‘Fonds voor Wetenschappelijk Onderzoek -- Vlaanderen’, G066215N, G0D5117N and G0B9317N). GB acknowledges support from the Interne Fondsen KU Leuven / Internal Funds KU Leuven under grant agreement C14/18/094, and the Research Foundation – Flanders (‘Fonds voor Wetenschappelijk Onderzoek – Vlaanderen’, G0E1420N).

## Supplementary material

**Table S1**. Accession numbers and GISAID labels for sequences used here. Note that EPI_ISL_406592, EPI_ISL_406595, EPI_ISL_403931, and EPI_ISL_402120 were excluded from our phylogenetic analyses.

**Table S2**. Log marginal likelihoods estimated for all analyses. The labels match those in Figure 1.

